# Endothelial cells filopodia participation in the anastomosis of CNS capillaries

**DOI:** 10.1101/415307

**Authors:** Miguel Marin-Padilla, Louisa Howard

**Affiliations:** Geisel School of Medicine, Dartmouth, College, Hanover, New Hampshire, 03755, USA

**Keywords:** Anastomosis, CNS capillaries, endothelial cell flopodia, flopodia conglomerates, post-anastomotic new CNS capillaries

## Abstract

By combining the classic Golgi method and the electron microscope, we have gained a better understanding of the anastomosis of CNS blood capillaries. The participation of growing capillary’ leading endothelial cells filopodia in the anastomotic process is described. The two approaching capillaries leading endothelial cells filopodia intermingle and interact forming complex conglomerates with narrow spaces filled with proteinaceous material (possibly basal lamina) secreted by them. The presence of tight junctions among the filopodia corroborates their vascular nature. Their presence also suggests a different endothelial cells origin as will those from the two approaching capillaries. The original narrow spaces coalesce into larger ones leading to the eventual formation of a single one that will interconnect (anastomose) the two capillaries. The newly formed post-anastomotic CNS capillaries are rather small with irregular and narrow lumina that might permit the passage of fluid but not yet of blood cells. Eventually, the new capillaries lumina will enlarge permitting the passage of blood cells.

**Funding information:** The M. M-P. Golgi studies were supported by a “Jacob Javits Neurosciences Investigator Award". NIH Grant NS-22897. And the L. H. EM studies were supported by the Gilman Fund/Class of 1978 Life Sciences Center. M. M-P. is Emeritus Professor of Pathology and Pediatrics and L. H. is a Consulting Electron Microscopists. Both from the Geisel School of Medicine at Dartmouth. Hanover, NH 03755, USA

## 1 INTRODUCTION

Recent investigations, using the classic Golgi method and the transmission electron microscope, have disclosed some developmental aspects of the human cerebral cortex microvascular system unknown previously, including: a) the equidistant perforations (ca. 500 *µm*) of the cortex external glial limiting membrane by meningeal capillaries (Marín-Padilla, 2011, 2012); b) the establishment of a Virchow-Robin Compartment, open to the meningeal interstitium, around each perforating vessels (Marín-Padilla and Knopman, 2011); and, c) the establishment of separated microvascular systems for the gray (GM) and white (WM) matter respectively (Marín-Padilla, 2015). A continuous and rapid one for the GM, where most neurons reside and, a slower one for the WM, essentially deprived of neurons. In the GM, blood enters through arterial perforators and exit through contiguous venous ones, supplying oxygen continuously and rapidly to GM neurons. The WM blood is supplied by the extension of a few GM arterial perforators interconnected by an extensive anastomotic plexus of thin-wall venules. The GM blood drains through the venous perforators into arachnoid veins and eventually into the dural sinuses. The WM venules drains into the periventricular venous plexus and through the Rosenthal veins into the ventral venous circle and through the Galen ampulla into the dural sinuses.

During the GM developmental maturation innumerable contacts (anastomoses) between growing capillaries occur with the establishment of a common (shared) lumen for the passage of blood. Vascular anastomoses also occur during the development of any organ and/or tissue and play crucial roles in both traumatic and surgical wounds healing as well as in the repair of bone fractures. It represents one the most generalized biological process known. Despite its widespread incidence and biological importance, some aspects of capillary anastomoses are still unexplained.

Various mechanisms on capillary anastomoses have been proposed: the presence of additional endothelial cells with formation of multicellular tubes and extracellular spaces (Blum et al., 2008), the rearrangements of endothelial cells with formation of multicellular tubes (Herwig et al., 2011), the elongation of endothelial cells with reorganization of junctions at the growing capillaries head, the confluence and canalization of small fissures into larger ones as well as the presence of additional endothelial cells at the head of growing capillaries (Ortuño Pacheco, 1971). Although all of those cellular events do occur in the anastomotic process, how they contribute to it remains unclear. Recent studies using color videos in living animals have shown extraordinary and beautiful imagen of blood vessels approach, fusion and establishment of a common (shared) lumen for the passage of blood (Lenard et al., 2015). Despite these videos, clarity, beauty and accuracy, at the photographic magnification used, it may not be possible to visualize the leading endothelial cells filopodia of the approaching capillaries. Even, their possible role in the anastomotic process has been recently questioned with the safeguard they could participate in some types of tissues (Phng et al., 2013).

We are revisiting the role of the leading endothelial cells filopodia in the anastomosis of capillaries in the mouse developing brain.

**FIGURE 1.**
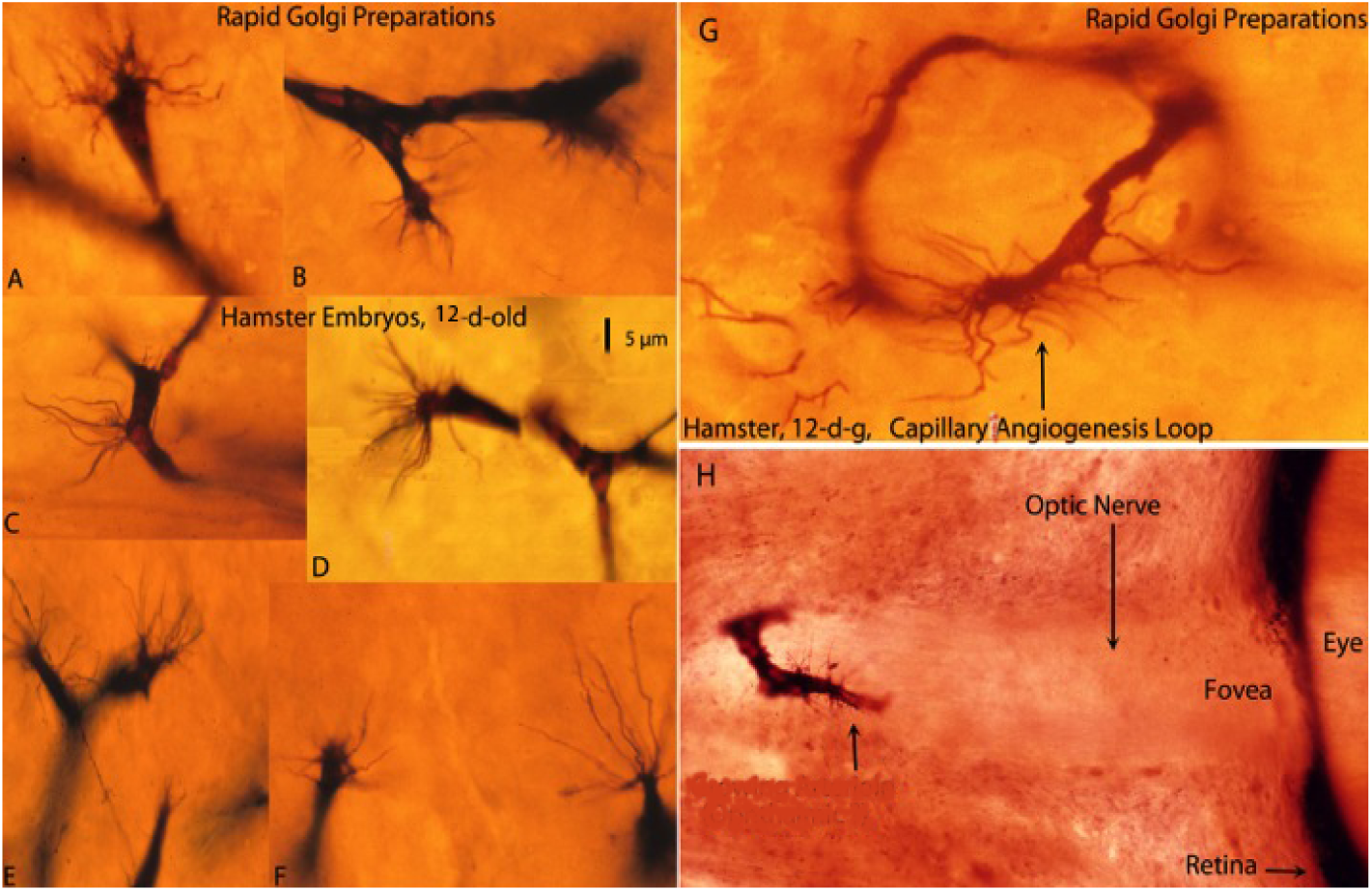
Composite figure of rapid Golgi preparations showing growing capillaries in the cerebral cortex of 12-d-o hamster embryos. Growing capillaries have a polypoid head with numerous radiating filopodia (A, B, D, E, and F). C. Detail of a new capillary growing directly from the vessel wall. G. Detail of a newly established capillary loop with some peripheral filopodia that have not participated in the anastomotic process. H. Detail of a growing capillary (arrow) with leading filopodia entering the optic nerve of a hamster embryo eye. The retina and the fovea are also shown. The 5 *µm* scale (D) is roughly applicable to all capillaries of the figure excepting the eye one (H). (from: Marín-Padilla (2012))

## 2 MATERIAL AND METHODS

In this study we have used the Golgi collection of one of us (M. M-P) that includes thousands of preparations of the embryonic cerebral cortex of hamsters, mice, cats and humans (Marín-Padilla, 2011). The classic Golgi method uses thick tissue preparations (150 to 170 *µm*), permitting three-dimensional views of growing CNS capillaries and of their radiating filopodia (Fig. 1). The various events in CNS microvascular anastomoses (approach, recognition, contacts and lumen sharing) and the role of leading endothelial cells’ filopodia are better envisaged in the Golgi thick preparations (Fig. 1). Once the main events in the anastomosis of CNS capillaries were better understood, they were subsequently explored using the transmission electron microscope (TEM). For the TEM studies, one of us (L.H.) made new recuts from the EM blocks of a previous developmental study of the mouse olfactory nerve that include the embryonic mouse cerebral cortex (Marin-Padilla and Amieva B., 1989). L.H. made 70 new thin sections (70 nm) from 11.5 day-old mouse embryos developing cortex. All TEM images were taken at 100 kV on a JEOL TEM1010 equipped with a XR-41B AMT digital camera and capture engine software (AMTV540; Advanced Microscopy Techniques).

**FIGURE 2.**
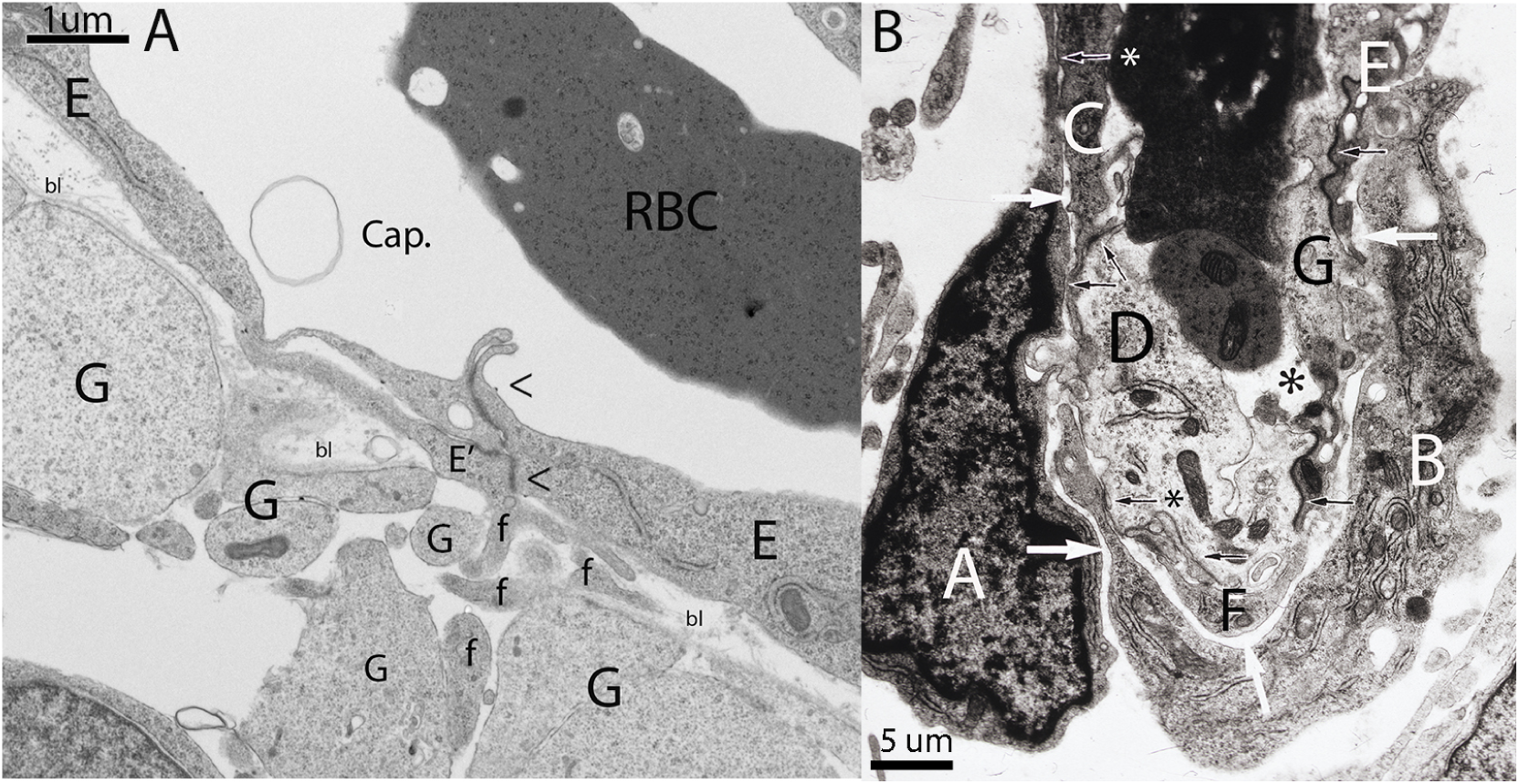
Composite figure of electron photomicrographs showing some aspects of a perforating pial capillary entering the cortex (A) and the advancing head (B) of a perforating meningeal capillary. A. Detail of a perforating meningeal capillary with an additional endothelial (E’) cell with advancing filopodia (f) that have perforated the cortex’ external glial limiting membrane (EGLM) composed of interconnected glial endfeet (G) covered by basal lamina (bl). Some of the filopodia (f) of the perforating EC have already enter into the nervous tissue. The perforating endothelial cell (E’) shares tight junction (<) with those of the capillary wall (E). Scale: 1 *µm*. B. View of the advancing polypoid head of a perforating meningeal capillary with nucleated red cells in its lumen (*), illustrating its ECs composition and organization. Some of the ECs (B, C, D, E, F, G) are components of the capillary wall and share tight junctions (black arrows). The two additional ECs (A and B) explain its bulging polypoid head (see Fig. 1). The additional ECs (A, B) are separated from those of the capillary wall by a long space (white arrows) but share small tight junctions with those of the capillary wall (arrows with asterisks). They are also the only ones with searching filopodia (see Fig. 1). Scale: 5 *µm*.

## 3 RESULTS

Microvascular anastomoses are complex processes that involve various sequential events, including: a) capillary approach and recognition; b) growing capillaries leading endothelial cells (ECs) filopodia contacts and integrations; c) establishment among the contacting filopodia of complex conglomerates with narrow unconnected accessory lumina (fissures) filled with proteinaceous material (possibly basal lamina) secreted by them; and, d) progressive confluence of the accessory lumina into larger ones leading to the establishment of a common one that eventually will interconnect (anastomose) the approaching capillaries allowing the circulation of blood. These events are accompanied by EC mitoses, addition of new ECs at the growing capillaries head and ECs displacements. The four additional ECs (two from each approaching capillaries) are not components of the capillary wall and are free to move. These additional ECs are the only ones with searching filopodia and are the mayor contributors in the anastomotic process (Figs. 1, 2).

### 3.1 Golgi studies

In Golgi preparations, CNS capillaries are stained black against a transparent yellowish background (Marín-Padilla, 2011). The advancing heads of growing capillaries are characterized by a polypoid head with numerous radiating filopodia (Fig. 1, A-G). The radiating filopodia of growing blood capillaries have been seldom described in the literature. Their polypoid head with radiating filopodia was originally described by Klosovskii, a Russian investigator, in a book with numerous illustrations (Klosovskii, 1963). They have been recently described in the embryonic brain by Puelles et al. (1976) and, in greater detail, in Marín-Padilla (2011, 2012, 2015) Golgi studies. It is possible that Cajal might have also recognized them but interpreted as glial filaments (Cajal, 1911).

New capillaries can emerge, as well as recede, from any place along the capillary wall (Fig. 1C). They establish anastomotic loops between different tissue labels (Fig. 1G). A growing capillary with advancing filopodia entering the optic nerve of a hamster embryo’ eye is also illustrated (Fig. 1H).

### 3.2 Electron-microscopic studies

Because, the Golgi study failed to recognize the ECs filopodia interactions that precede the anastomotic event, an electron-microscopic investigation was carried out. CNS blood capillaries are easily recognized in EM preparations of the developing cortex as hollow structures (5 to 7 *µm* in diameter) with nucleated red cells and walls composed of endothelial cells separated by tight junctions.

The EM studies show that the polypoid head of growing CNS capillaries is composed of a variety of endothelial cells (Fig. 2B). Some are components of the capillary wall separated by tight junctions (Fig. 2B: C, D, E and F). The growing capillary head has two additional ECs (Fig. 2A, B). They are separated from the capillary wall by long narrow spaces and are free to move (Fig. 2B, white arrows). They do share small tight junctions with those of the capillary wall corroborating their vascular nature (Fig. 2B, black arrows with asterisks). The four additional ECs (two from each approaching capillary) are the only ones with searching filopodia. The meningeal capillary that perforates the cortex external glial limiting membrane (EGLM) to enter into the brain also has an additional EC with advancing filopodia (Fig. 2A, f).

Four developmental stages in the anastomosis of CNS capillaries were explored in the TEM preparations, including: a) the complex interconnections (conglomerates) among the leading ECs filopodia from the approaching capillaries; b) the formation among the contacting filopodia of narrow unconnected spaces (accessory lumina) filled with proteinaceous material, possibly basal lamina, secreted by them; c) confluence of accessory lumina into larger ones, forgoing the common one that will interconnect (anastomose) the approaching capillaries; and, d) the newly formed and very small post-anastomotic CNS capillaries.

The interactions of the leading ECs filopodia from the approaching capillaries represent a fundamental event in the anastomosis of blood vessels in the developing brain. The leading ECs filopodia from the two approaching capillaries establish complex interactions (conglomerates) that are difficult to be recognized as vascular structures (Fig. 3A). Their morphology and organization vary significantly depending on their developmental stage and the TEM section angle. Cross sections of these filopodia conglomerates, as the one illustrated in Fig. 3A, are the most illustrative. To identify them as vascular structures they will need to have tight junctions among the contacting filopodia (Fig.3A, small black arrows). Their presence among the contacting filopodia suggests different endothelial cells origins, as will be those from the two approaching capillaries ECs. Some of the filopodia conglomerates might even have a section of the nucleus of one of the participant endothelial cells, further corroborating their vascular nature (Fig. 3A, N). It is proposed that confluence of small lumina into larger ones embodies the eventual formation of a single one that will anastomose the two approaching capillaries allowing the circulation of blood between them. The continuous secretion of basal lamina material by the filopodia could play a role in the confluence of the accessory lumina.

The filopodia conglomerates have also additional peripheral filopodia that have not participated in the anastomotic process (Fig. 3A). Residual non-participating peripheral filopodia are also recognized around the newly formed post-anastomotic CNS capillaries (Figs. 3B, C, D) as well as around recent anastomosis (Fig. 1G). Once the anastomotic process is completed the non-participating filopodia are reabsorbed.

In the EM preparations studied there were also minute capillaries composed of at least two endothelial cells separated by tight junctions and very small, narrow and irregular lumen (Fig. 3B, C, D). We interpreted these small capillaries as newly formed post-anastomotic CNS capillaries. Some of them may still has more than one lumen reflecting their multiple origin (Fig. 3B, asterisks). Others have very small, narrow and irregular lumen (Fig. 3C, asterisk). Others have narrow and convoluted lumen (Fig. 3D, asterisks). At this developmental time, the newly formed post-anastomotic CNS capillaries might permit the passage of fluids but not yet of blood cells. Eventually, they will enlarge permitting the passage of blood cells.

## 4 DISCUSSION

The CNS growing capillaries’ leading ECs filopodia functions, include: the control of the capillary directional growth, the recognition of those of other growing capillaries and the establishment of complex interconnections (conglomerates) with them prior to their anastomosis.

To visualize the ECs filopodia of growing CNS capillaries number, size and radiation requires the use of special techniques that utilizes thick preparations, such as the classic Golgi method. The rare usage of his classic and old method (Golgi, 1873) might explain why the ECs filopodia of growing capillaries has been so seldom described in the literature. Even at the photographic magnification used in the extraordinary color videos of vascular anastomosis in the living animal will not permit to visualize them.

Microvascular anastomoses are complex processes that involve various sequential events, including: a) capillary approach and recognition; b) the approaching capillaries leading endothelial cells (ECs) filopodia contacts and integrations; c) the establishment among the contacting filopodia of complex aggregates (conglomerates) with narrow unconnected accessory lumina (fissures) filled with proteinaceous material (possibly basal lamina) secreted by them; and, d) the progressive confluence of the accessory lumina into larger ones leading to the establishment of a common one that eventually interconnect (anastomose) the approaching capillaries allowing the circulation of blood. These events are accompanied by EC mitoses and by the addition of new ECs at the head of growing capillaries. The four additional ECs (two from each approaching capillaries) are not actual components of the capillary wall and are free to move. They are also the only ECs with searching filopodia and mayor contributors in the anastomotic process (Figs. 1, 2).

**FIGURE 3.**
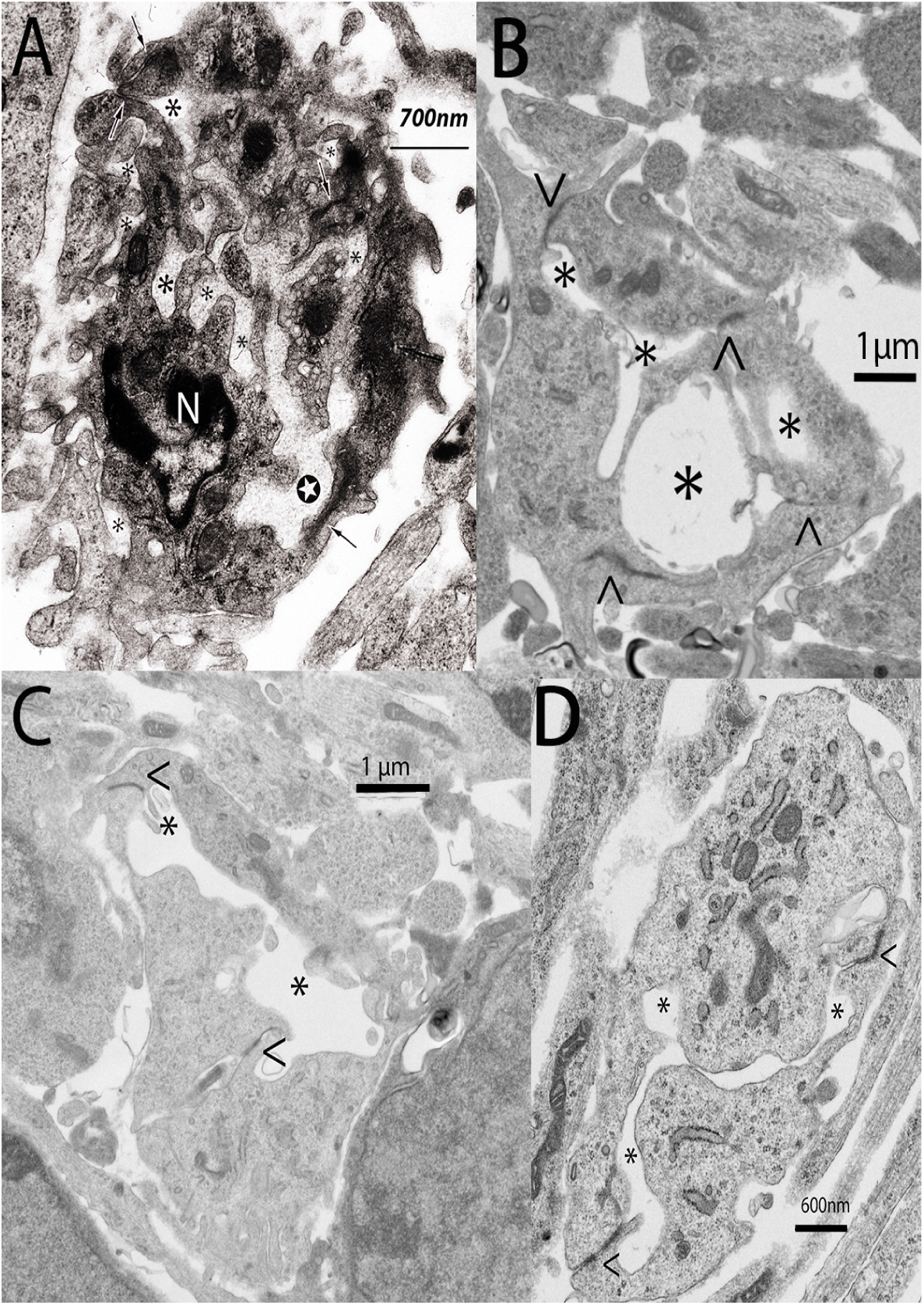
Montage of electron photomicrographs, including a complex conglomerate of EC filopodia (A) from the approaching CNS capillaries and recently formed and rather small (B, C, D) post-anastomotic CNS capillaries. A. Detail of a complex filopodia’ conglomerate with narrow spaces or fissures (asterisks) filled with proteinaceous material (possibly basal lamina) produced by them. The morphology and organization of these filopodia conglomerates vary considerably and depend on the angle of the section and its developmental stage. It is often difficult to recognize them as vascular structures. The presence of tight junctions among the filopodia (arrows) confirm their vascular nature and suggest a different endothelial cells origin, as will be those from the two approaching capillaries. Some even have a section of the nucleus (N) from one of the approaching capillaries endothelial cell. Often one of the lumina (white star) is larger than the rest suggesting the confluence of smaller ones. B. View of a new post-anastomotic CNS capillary composed of three endothelial cells separated by tight junction (<, >) with three irregular lumina (asterisks), reflecting its multiple luminal origin. C, D. Views of recently formed new post-anastomotic CNS capillaries composed of two endothelial cells separated by tight junction (<, >) with very small and narrow single lumen (asterisks). At this stage, the post-anastomotic new CNS capillaries will permit the passage of fluids but not yet of blood cells. Eventually, their lumen will enlarge and permit the passages of blood cells. The new post-anastomotic CNS capillaries (B, C, D) still have some peripheral filopodia that have not participated in the anastomotic event. Scales: 3A (700 nm), 3B (1 *µm*), 3C (1 *µm*) and 3D (600 nm).

The polypoid head of growing blood capillaries has two additional endothelial cells separated from those of its wall by a long narrow space but sharing small tight junctions with them (Figure, 2B). Their functions include the recognition of those of other growing capillaries and the establishment of complex aggregates (conglomerates) with them. The resulting conglomerates are complex structures composed of numerous filopodia separated by narrow spaces filled with proteinaceous material, possibly basal lamina, segregated by them. Their vascular nature is corroborated by the presence of tight junctions among the contacting filopodia. Their presence suggests different endothelial cell origin, such as those from the two approaching capillaries. Some filopodia conglomerates may have a section of the nucleus from one of the approaching capillaries ECs, further corroborating their vascular nature.

We propose that the original spaces (fissures) formed among the leading ECs filopodia will coalesce into a single one that interconnect (anastomose) the approaching capillaries allowing the passage of blood. The progressive accumulation of basal lamina material among the filopodia may contribute to their fusion. The four leading ECs (two from each approaching capillary) become the wall of the newly formed post-anastomotic CNS capillaries. They are components of the approaching capillaries and are attached to them by small tight junctions. How the lumen of the two approaching capillaries interconnect to the single one at the center of the filopodia conglomerates need to be further investigated.

The newly formed post-anastomotic CNS capillaries are rather small, ranging between 2.5 and 3.5 *µm* in diameter. Some of them might still have more than one lumen (reflecting their multiple origin) and/or a very narrow and single ones. At this stage, the new post-anastomotic CNS capillaries might permit the passage of fluid but not yet of blood cells. Eventually, their lumen will enlarge and permit the passage of blood cells.

There are no reasons to think that microvascular anastomosis in other developing tissues will not establish similar filopodia conglomerates between their approaching capillaries. Microvascular anastomosis in different developing tissues need to be further investigated with appropriate methods.

## Appendix

